# Preparation of Synaptosomes from Postmortem Human Prefrontal Cortex

**DOI:** 10.1101/006510

**Authors:** Neil R. Smalheiser, Giovanni Lugli

## Abstract

Synaptosomes are a popular type of isolated synaptic fraction intensively used in neuroscience and cell biology. They are prepared by layering on density gradients and thought to consist largely of axonal endings with attached postsynaptic structures (Morgan, 1976), in contrast to synaptoneurosomes (Hollingsworth et al, 1985) which are prepared by filtration and are thought to consist largely of pinched-off dendritic spines with attached presynaptic structures. Although most studies of synaptosomes have utilized rodent or primate tissue, a score of studies have employed human samples derived from surgical specimens or postmortem brain. We recently described the isolation of synaptosomes from human postmortem prefrontal cortex to study the expression of synaptic microRNAs and other small RNAs in depression, schizophrenia and bipolar disorder (Smalheiser et al, 2014). This protocol alluded to methods and modifications that are scattered among several publications, and did not explain the reasons for the procedures chosen. Because our protocol differs from other published human synaptosome protocols in a variety of respects, we present here a detailed description of synaptosome preparation that should facilitate the use of this standardized synaptic fraction by other workers.

## Materials and Methods

1. Tissue dissection. Stored as frozen unfixed slices at −80 degrees. Samples (50-100 mg wet weight per prep) are taken that span across all layers (including white matter as well as grey matter), and are minced at −80.
2. Tissue homogenization. Each minced prep is immediately homogenized using a Dounce pestle in ice-cold homogenization buffer (HB) containing a cocktail of protease and RNase inhibitors [50 mM Hepes, pH 7.5, 125 mM NaCl, 100 mM sucrose, 2 mM K acetate, 10 mM EDTA, 2 mM phenylmethylsulfonyl fluoride, 10 mM N-ethylmaleimide, 10 μg/mL leupeptin, 1 μg/mL pepstatin A, 2 μg/mL aprotinin, 160 U/mL Superase-In (Ambion), 160 U/mL Rnase-OUT (Life Technologies, Carlsbad, CA, USA)]. Aliquots of this “total tissue homogenate” are saved for protein or RNA isolation, to ensure molecular integrity of the sample and as a control for synaptic enrichment of the synaptosomes.
3. Centrifugation. After discarding material that pellets at low speed (1,000 x g for 10 minutes at 4 degrees), a crude membrane pellet ‘P2’ is obtained by centrifugation at 20,000×g for 20 min. (Note: For some purposes, the P2 fraction may be utilized as a poor-man’s proxy for a more purified synaptic fraction, particularly if the amount of available tissue is limiting.)
4. Layering on discontinuous sucrose gradient. The P2 pellet is resuspended in 0.32 M sucrose in 1 mM NaHCO_3,_ layered on a discontinuous gradient of 0.32, 0.85, 1.0, and 1.2 M sucrose in 1 mM NaHCO_3_, and centrifuged at 100,000×g for 2 h. The synaptosomal fraction is recovered as a band at the 1.0–1.2 M interface. The synaptosomes are quickly pelleted at 20,000×g for 20 min and rinsed twice in 4 × volume of homogenization buffer containing protease inhibitors and spun down again 20,000×g for 20 minutes prior to extracting RNA. Note that the sucrose is treated with RNAsecure (Ambion) as per the manufacturer’s instructions before making the sucrose gradient.

## Results and Discussion

From 50 mg wet weight of prefrontal cortex, we generally obtain ∼2 micrograms of total RNA from isolated synaptosomes. Like synaptosomes prepared from fresh mouse brain tissue, the human synaptosomes appear to have relatively intact RNA integrity as shown by RNA bioanalyzer profiling and gel electrophoresis, and show the expected enrichment in synaptic proteins and RNAs and depletion of nuclear markers (Smalheiser et al, 2014).

It should be noted that some workers have employed homogenization buffers containing iso-osmotic sucrose (0.32 M) with low salt (Cohen et al, 1977). Use of iso-osmotic conditions is intended to minimize swelling, shrinkage or rupture of organelles. We have chosen instead to reduce sucrose (to 0.1 M) and to maintain physiological ionic strength by adding salts, which should minimize nonspecific protein binding to synaptosomes.

One should consider whether to include Ca^2+^ and Mg^2+^ during homogenization. We chose not to (and in fact, add calcium chelators), in order to avoid activating RNAses. Additional RNAse inhibitors were added to further reduce the risk of altering RNAs in the preparation. However, if one wishes to keep RNA-protein complexes intact, the inclusion of Mg^2+^ is advisable. Note, also, that polyribosomes are best kept intact by treating samples with cycloheximide (e.g., Stefani et al, 2004).

Our group has not prepared PSDs from human material, though we would not anticipate any difficulty in doing so. To obtain isolated postsynaptic densities (PSDs), the classic protocol is to subject synaptosomes to hypotonic lysis followed by Triton X-100 or NP-40 detergent extraction of the pellet (Morgan, 1976; Cohen et al, 1977). However, if one is simply interested in the PSD and not in the material released by lysis (including synaptic vesicles and other pre- and postsynaptic cytoplasmic elements), one might skip the osmotic lysis step and carry out detergent extraction.

## Acknowledgements

Our studies were supported by the Stanley Medical Research Institute and the Alzheimer’s Association.

